# Integration of Brassinosteroid and Phytosulfokine Signalling Controls Vascular Cell Fate in the *Arabidopsis* Root

**DOI:** 10.1101/244749

**Authors:** Eleonore Holzwart, Apolonio Ignacio Huerta, Nina Glöckner, Borja Garnelo Gómez, Friederike Ladwig, Sebastian Augustin, Jana Christin Askani, Ann-Kathrin Schürholz, Klaus Harter, Sebastian Wolf

**Affiliations:** Department of Cell Biology, Centre for Organismal Studies Heidelberg, Heidelberg University, Im Neuenheimer Feld 230, 69120 Heidelberg, Germany; Plant Physiology, ZMBP, Universität Tübingen, Auf der Morgenstelle 32, 72076 Tübingen, Germany

**Author notes:** Present address: Department of Biology, ETH Zurich, Universitätstrasse 2, 8092 Zurich, Switzerland. Present address: Department of Plant Molecular Biology, University of Lausanne, 1015 Lausanne, Switzerland (Sebastian Augustin). Corresponding author: Sebastian Wolf, Im Neuenheimer Feld 230, 69120 Heidelberg, Germany. **Author contributions:** EH, AIH, NG, BGG, FL, SA, JCA, and A-KS performed experiments and analysed data. KH designed research and analysed data. SW performed experiments, designed research, analysed data and wrote the manuscript.

**Keywords:** cell fate, xylem, procambium, brassinosteroids, phytosulfokine, signalling, cell wall, development, differentiation, root, Arabidopsis

## Abstract

Multicellularity arose independently in plants and animals, but invariably requires robust determination and maintenance of cell fate. This is exemplified by the highly specialized water-and nutrient-conducting cells of the plant vasculature, which are specified long before their commitment to terminal differentiation. Here, we show that the hormone receptor BRASSINOSTEROID INSENSITIVE 1 (BRI1) is required for root vascular cell fate maintenance, as BRI1 mutants show ectopic xylem in procambial position. However, this phenotype is unrelated to classical brassinosteroid signalling outputs. Instead, BRI1 is required for the expression and function of its interaction partner RECEPTOR-LIKE PROTEIN 44 (RLP44), which, in turn, associates with the receptor for the peptide hormone phytosulfokine (PSK). We show that PSK signalling is required for the maintenance of procambial cell identity and is quantitatively controlled by RLP44, which promotes complex formation between the receptor for PSK and its co-receptor. Mimicking the loss of RLP44, PSK-related mutants show ectopic xylem in the position of procambium, whereas *rlp44* can be rescued by exogenous PSK. Based on these findings, we propose that RLP44 controls cell fate by connecting BRI1 and PSK signalling, providing a mechanistic framework for the integration of signalling mediated by the plethora of plant receptor-like kinases at the plasma membrane.

## Introduction

A key function of signalling networks in multicellular organisms is to ensure robust determination and maintenance of cell fate. In plants, extreme specialization is displayed by the cells of the vascular tissues, vital for the distribution of water, nutrients, and signalling molecules. Xylem tracheary elements are characterized by lignified secondary cell wall thickenings that protect against collapse and provide mechanical support for vertical growth. Positioned between xylem and the nutrient-transporting phloem are the cells of the procambium, which give rise to the lateral meristems during secondary growth (1). Root vascular tissue patterning is set up in the embryo and maintained in the post-embryonic root by mutual antagonism of auxin and cytokinin signalling domains (2-5). After xylem precursor cells are displaced from the root meristem, an intricate gene-regulatory network connected to patterning mechanisms by HD-ZIP III transcription factors mediates differentiation into tracheary elements (6-9). Thus, primary root xylem cell fate can be traced back to early specification events in the embryo. In contrast, during secondary growth, (pro)cambial cells adjacent to the existing tracheary elements acquire xylem cell fate dependent on positional information (10).

Brassinosteroid (BR) hormone signalling (11, 12) has been implicated in xylem differentiation and vascular patterning (13, 14). BRs are perceived by BRASSINOSTEROID INSENSITIVE 1 (BRI1) (15) which belongs to the large group of plant receptor-like kinases (RLK) with a leucine-rich repeat (LRR) extracellular domain, a transmembrane domain, and a cytosolic kinase domain related to animal Irak and Pelle kinases (16). Upon ligand binding, BRI1 heterodimerizes with members of the SOMATIC EMBRYGENESIS RECEPTOR KINASE (SERK) LRR-RLK family such as BRI1 -ASSOCIATED KINASE 1 (BAK1)(17, 18), and activates a signalling cascade that negatively regulates BRASSINOSTEROID INSENSITIVE 2 (BIN2)(19), a GSK3-like kinase that phosphorylates the BR-responsive transcription factors BRASSINAZOLE RESISTANT 1 (BZR1)(20) and BRI1 EMS SUPPRESSOR 1 (BES1)/BZR2. Inhibition of BIN2 activity allows BZR1 and BES1 to translocate to the nucleus, where they mediate BR-responsive control of transcription (21-23). A so far somewhat enigmatic relationship exists between BR and PHYTOSULFOKINE (PSK) signalling. PSKs are small secreted peptide growth factors that have been implicated in a variety of fundamental processes and are perceived by two close relatives of BRI1, PHYTOSULFOKINE RECEPTOR 1 and 2 (24-27). PSK activity depends on proteolytic processing of the precursor peptides, the sulfation of two tyrosine residues in the mature pentapeptide (YIYTQ) by TYROSYLPROTEIN SULFOTRANSFERASE (TPST) (28), and functional BR signalling (29-31). At present, it is not clear how BR and PSK signalling interact but the receptors for both growth factors share the requirement for a SERK co-receptor (32, 33).

Recently, we demonstrated that feedback information from the cell wall is integrated with BR signalling at the level of the receptor complex through RECEPTOR-LIKE PROTEIN (RLP) 44 (34). RLP44 is genetically required for the BR-mediated response to impaired cell wall modification and is sufficient to elevate BR signalling when overexpressed. RLP44 was shown to be in a complex with BRI1 and BAK1 and to directly interact with BAK1. Thus, we hypothesized that RLP44 modulates BR signalling strength in response to cues from the cell wall (34). However, it is not clear whether the RLP44-BR signalling module plays additional roles in plant physiology, besides a possible mechanism for cell wall homeostasis (35). Here, we show that RLP44 is required for the maintenance of cell fate in the root vasculature by connecting components of the BR and PSK signalling pathways. RLP44 directly interacts with BRI1 and controls xylem differentiation in a BRI1-dependent manner, but independently of BR signalling outputs. In addition, RLP44 can directly interact with PSKR1, and the *rlp44* phenotype can be rescued by application of the PSK peptide. Moreover, mutants affected in PSK signalling show an rlp44-like xylem phenotype, suggesting that RLP44 has a positive effect on PSK signalling, which, in turn, promotes procambial identity.

## Results

### RLP44 directly interacts with the brassinosteroid receptor BRI1

We previously demonstrated that RLP44 is present in a complex with both BRI1 and its co-receptor BAK1 and is able to promote BR signalling upon cues from the cell wall or when overexpressed. In addition, we provided evidence for direct interaction with BAK1 (34). To assess whether RLP44 can also directly interact with BRI1, we performed a mating-based split ubiquitin assay in yeast (36). Under selective conditions, interaction of BRI1 and RLP44 enabled yeast growth, similar to what was observed before with BAK1-RLP44 and BAK1-BRI1 (34) (Fig. 1A). In addition, Foerster resonance energy transfer-fluorescence lifetime imaging microscopy (FRET-FLIM) analysis after transient expression in *Nicotiana benthamiana* leaves confirmed that BRI1 and RLP44 are able to interact directly (Fig. 1B and Fig. S1A). Furthermore, endogenous BRI1 and BAK1 were detected in immunoprecipitates of *RLP44-GFP* expressed under the control of its own promoter in the *rlp44^cnu2^* mutant background (Fig. S1B). In summary, RLP44 and BRI1 form complexes in yeast and *in planta* through a direct interaction. To assess the potential role of the RLP44-BRI1 signalling modules we sought to identify the tissues in which it might be active. As BRI1 is ubiquitously expressed, we thus investigated the expression pattern of RLP44.

**Fig. 1.**
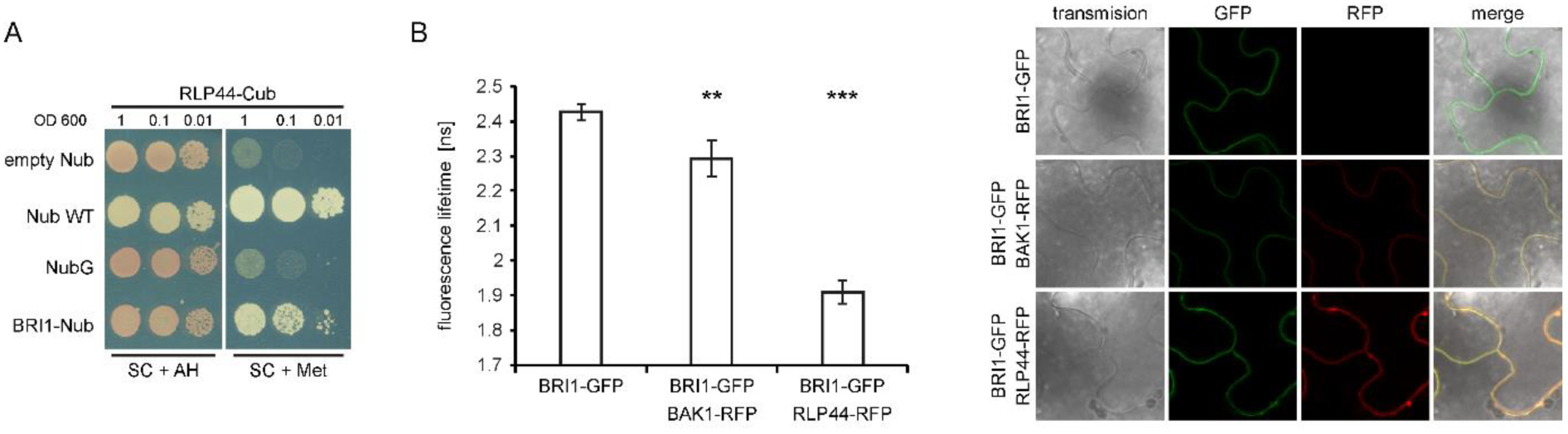
RLP44 directly interacts with BRI1. (A) Mating-based split-Ubiquitin analysis in yeast reveals direct interaction of RLP44 and BRI1. After mating, presence of RLP44-Cub and BRI1-Nub enables yeast growth under selective conditions (SC + Met). Empty Nub vector and NubG are used as negative controls whereas interaction with WT Nub serves as a positive control. (B) FRET-FLIM analysis of the RLP44-BRI1 interaction in *Nicotiana benthamiana* leaves. Bars denote average of 5 measurements ± SD. Asterisks indicate statistically significant difference from BRI1-GFP after pairwise t-test (**p < 0.01; ***p < 0.001).

### RLP44 is expressed in the developing root vasculature

To study the function of RLP44, we generated transgenic plants expressing a translational GFP fusion of RLP44 under control of the RLP44 promoter (*pRLP44:RLP44-GFP*). These plants displayed elongated, narrow leaf blades and elongated petioles, reminiscent of BRI1 overexpressing plants (Fig. 2A and B) (37), and as previously observed with RLP44 overexpression (34). We crossed a *pRLP44:RLP44-GFP* line with the *RLP44* loss-of-function mutant *rlp44^cnu2^* derived from *comfortably numb 2* (34), which resulted in plants with a wildtype-like appearance (Fig. 1C), demonstrating that the fusion protein is functional, and confirming that the additional transgenic RLP44 expression was causative for the observed morphological effects (Fig. S2A). In the root apical meristem of *pRLP44:RLP44-GFP* and *pRLP44:RLP44-GFP* (*rlp44^cnu2^*), fluorescence was present in most tissues, but markedly enriched in epidermis and lateral root cap (Fig. 2D-F and Fig. S2B-E); slightly enhanced expression was also observed in xylem precursor cells (Fig. S2B). A strong increase of GFP fluorescence in the stele was observed towards the more mature part of the root (Fig. 2D-G and Fig. S2B-D), in accordance with previously published transcriptome data (38) and β-glucuronidase reporter activity under control of the RLP44 promoter (Fig. S2F, G). In the differentiating part of the root stele, RLP44-GFP fluorescence was relatively weak in the phloem, intermediate in xylem, and highest in the undifferentiated procambial cells (Fig. 1H, I, and Fig. S1C and D, compare Fig. 3A for vascular anatomy).

**Fig. 2.**
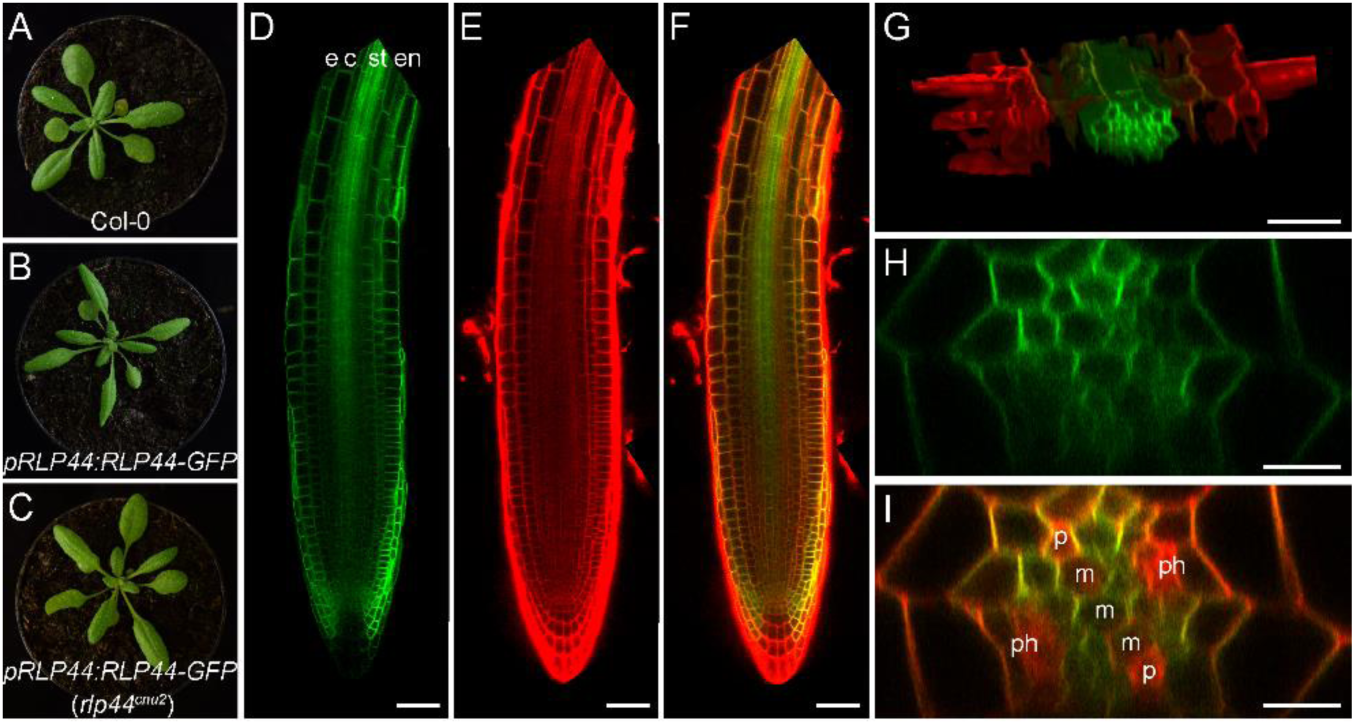
RLP44 is expressed in the root vascular tissue. (A-C) Effect of expressing *RLP44* as a translational fusion to GFP under control of its native 5’ regulatory sequence (*pRLP44:RLP44-GFP*). (A) Col-0. (B) *pRLP44:RLP44-GFP* in wildtype background shows a growth phenotype reminiscent of enhanced BR signalling. (C) Mutation of endogenous RLP44 in *pRLP44:RLP44-GFP* (*rlp44^cnu2^*) reconstitutes wildtype-like phenotype. (D-F) *pRLP44:RLP44-GFP* expression (D) in root meristem counterstained with propidium iodide (E) and merged (F). e = epidermis, c = cortex, st = stele, en = endodermis. Scale bars = 100 μm. (G) Projection of a confocal stack through the differentiation zone of a *pRLP44:RLP44-GFP* root showing fluorescence predominantly in the stele. (H and I) Optical section through the stele of a *pRLP44:RLP44-GFP* expressing root in the differentiation zone (H), counterstained with propidium iodide (I) indicating differentiated phloem (ph) and protoxylem (p) as well as yet undifferentiated metaxylem (m). Scale bar = 10 μm.

**Fig. 3.**
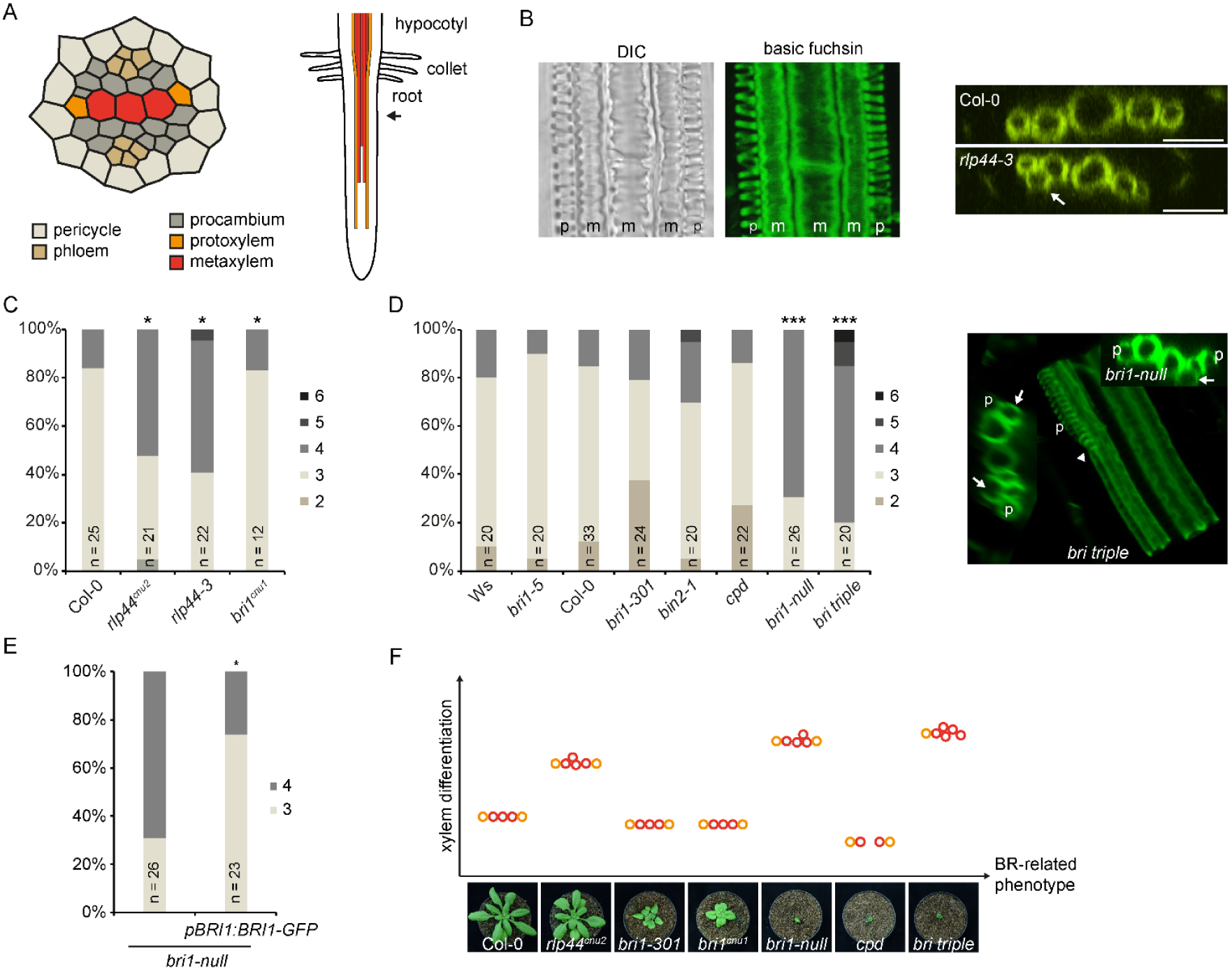
RLP44 and BR11 are required for the control of xylem cell fate. (A) Overview over xylem differentiation in the Arabidopsis root and schematic representation of the stele. Arrow marks point of xylem observation. (B) Basic fuchsin staining of 6 day old Arabidopsis root. DIC image shows secondary cell wall thickenings of protoxylem and metaxylem (left panel), basic fuchsin labels lignified secondary cell walls (middle panel). Confocal stacks allow xylem number quantification of the indicated genotypes in orthogonal view (right panel). Note ectopic metaxylem in procambial position (arrows). Left panel is a median plane image, middle panel is a maximum projection of a confocal stack. (C and D) Frequency of roots with the indicated number of metaxylem cells in *rlp44* and *bri1* mutants (C) and BR-related mutants (D) after basic fuchsin staining as in (B). Right panel in (D) shows orthogonal view and maximum projection of a confocal stack obtained with fuchsin stained *bri triple* mutant. Note metaxylem in procambial position (arrows) and disrupted protoxylem (arrowhead). Asterisks indicate statistically significant difference from Col-0 based on Dunn’s post-hoc test with Benjamini-Hochberg correction after Kruskal-Wallis modified U-test (*p < 0.05; ***p < 0.001). (E) Transgenic expression of *BRI1* under control of its own regulatory 5’ sequence rescues the ectopic xylem phenotype of *bri1-null*. (F) Overview over ectopic xylem phenotypes of *rlp44* and BR-related mutants. Note the absence of correlation between severity of BR signalling deficiency (x-axis) and frequency of ectopic xylem phenotype (y-axis). Based on (C) and (D).

### RLP44 controls xylem cell fate in a BRI1-dependent manner, but independently of BR signalling outputs

The prevalence of *pRLP44:RLP44-GFP* fluorescence in the stele prompted us to study the role of RLP44 in vascular development. To this end, we visualized lignified secondary cell walls in *rlp44* loss-of-function mutants through basic fuchsin staining and observed supernumerary metaxylem-like cells, frequently outside the primary xylem axis in the position of the procambium (Fig. 3A and B), a phenomenon we never observed in wildtype roots. Quantification of metaxylem cells in seedling roots of both *rlp44^cnu2^* and the T-DNA insertion line *rlp44-3* six days after germination (dag) showed a significant increase (Fig. 3C), suggesting that RLP44 controls xylem cell fate. Expression of RLP44 under control of its own promoter complemented this phenotype (Fig. S3). Since we had previously identified RLP44 as an activator of BR signalling, we also included *bri1^cnu1^*, a hypomorphic *bri1* allele (39) in the analysis. Surprisingly, primary root xylem number in *bri1^cnu1^* was indistinguishable from wildtype, suggesting that the ectopic xylem in *rlp44* mutants might be unrelated to the outputs of BR signalling. We therefore analysed the root xylem of a number of BR-related mutants spanning a broad range of growth phenotypes. Hypomorphic *bri1* mutants such as *bri1-301* (19) and *bri1-5* (40), the more severe signalling mutant *bin2-1* (19), and the BR-deficient biosynthetic mutant *constitutive photomorphogenic dwarf* (*cpd*) (41) did not show a pronounced increase in xylem cell number (Fig. 3D). In sharp contrast, *bri1* null alleles such as a previously characterized T-DNA mutant (termed *bri1-null*) (42) and the *bri1 brl1 brl3* triple mutant (called *bri triple* from hereon) (43) displayed a marked increase in the number of differentiated xylem cells (Fig. 3D), whereas expression of BRI1 under the control of its own promoter in *bri1-null* resulted in wildtype-like xylem (Figure 3E). Taken together, our results show that the xylem differentiation phenotype does not correlate with the severity of BR deficiency-related growth phenotypes (Fig. 3F). This is best exemplified by the comparison between the *cpd* and *bri1* null mutants, with *cpd* displaying wildtype-like or even slightly decreased xylem cell numbers, despite exhibiting a *bri1-*null-like growth phenotype. Thus, the control of xylem cell number, is largely independent of BR signalling outputs, but requires the presence of both BRI1 and RLP44. Interestingly, the expression of *RLP44* was reduced in the *bri1-null* mutant but not in *bri1* hypomorphs or *cpd*, suggesting that reduced RLP44 levels could partially explain the xylem phenotype of *bri1-null* (Fig. S4A, B). Consistent with this, uncoupling RLP44 transcription from BRI1 control through the 35S promoter could alleviate the *bri1-null* xylem phenotype (Fig. S4C), suggesting that BRI1 and RLP44 indeed act in the same pathway regulating xylem cell fate. In contrast, overexpression of *RLP44* had no effect on growth BL-insensitivity of *bri1-null* (Fig. S5). Taken together, our findings demonstrate that the phenotype of *bri1* loss-of-function mutants is partially independent from BR signalling outputs and suggest that RLP44 exerts its function downstream of BRI1 through other, yet unidentified signalling components.

It has been previously reported that root vascular development is responsive to environmental conditions (7). We compared xylem cell numbers of plants grown on standard medium (0.9% agar) and those that experienced increased mechanical force by growth on reclined hard agar plates (2% agar) (44). Interestingly, the number of xylem cells in the wildtype increased substantially under those conditions, whereas it did not further increase in *rlp44-3* (Fig. S6).

### Vascular cell fate determination by RLP44 and BRI1 is independent of BR signalling-mediated control of cell proliferation

We next asked whether the increase in xylem cell number observed in the *rlp44* mutant is reflected in enhanced cell proliferation in the root and therefore quantified vascular cell number of the seedling roots 6 dag. In *rlp44-3*, vascular cell number was indistinguishable from wildtype in the differentiation zone, suggesting normal cell proliferation in the root meristem (Fig. 4A). The *bri1^cnu1^* mutant, which did not display ectopic xylem cells, showed a significant increase in total vascular cell number (Fig. 4A), consistent with the recently described role of BR signalling in controlling formative cell divisions (45). These results suggest that increased xylem and increased proliferation in the vasculature are independent phenomena, and confirm a role for BR signalling in controlling formative cell divisions in the root meristem (Fig. 4A). In line with this, depletion of BRs in wildtype roots by application of the BR biosynthesis inhibitor PPZ (46) resulted in pronounced increase of vascular cell number (Fig. 4B, C). When PPZ-treated roots were supplemented with a low dose (0.5 nM) of BL, both root growth and vascular cell number were fully recovered (Fig. 4B, C). A higher-than-optimal dose of BL (5 nM) suppressed root growth and led to a strongly decreased vascular cell number (Fig. 4C). The *rlp44^cnu2^* mutant displayed a wildtype-like response to the manipulation of BR levels in terms of cell number (Fig. 4C), further supporting the independence of the xylem cell fate phenotype from the BR-signalling mediated control of cell proliferation (45).

**Fig. 4.**
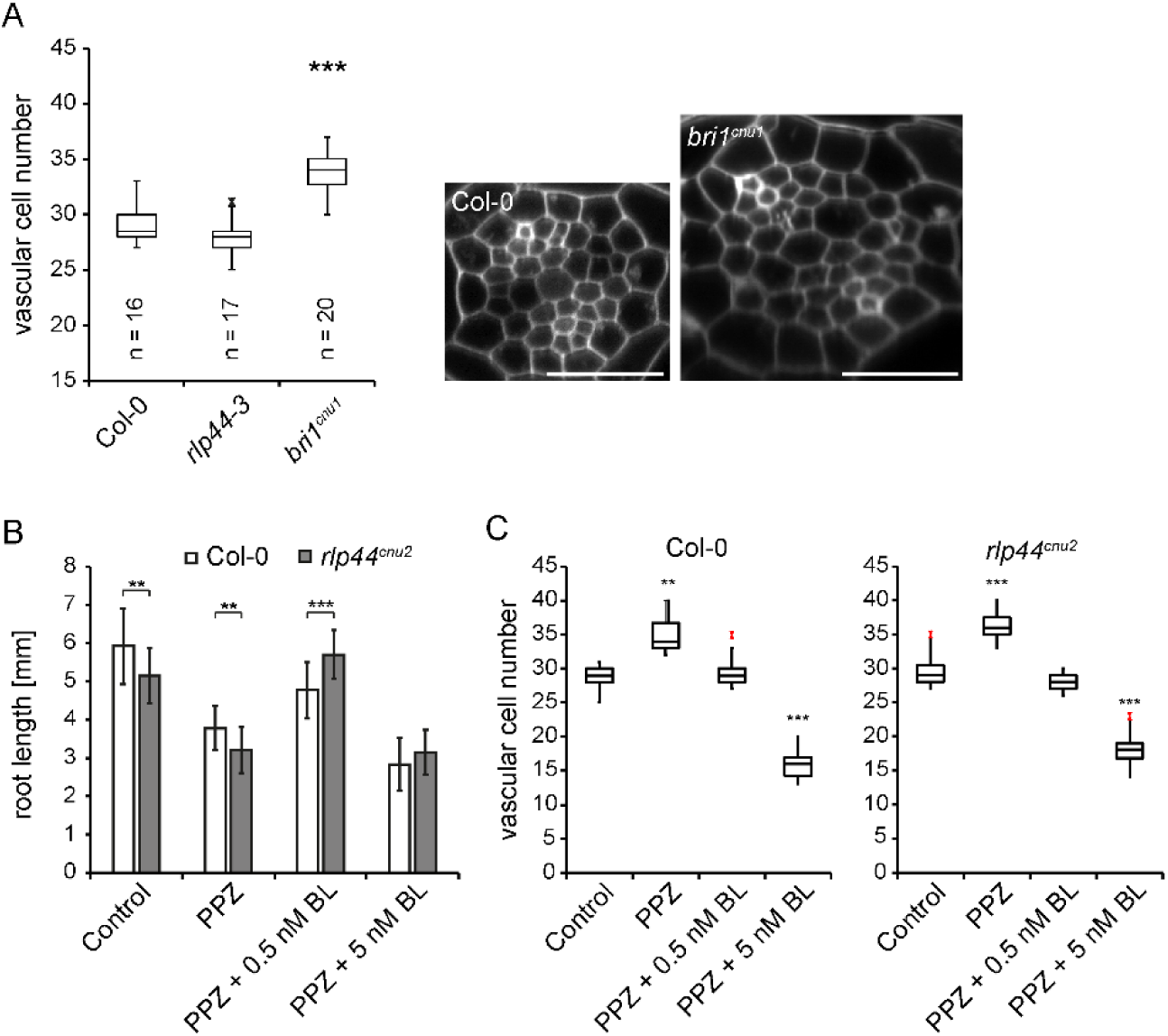
RLP44 and BRI1 determine vascular cell fate independent of BR signalling-mediated control of cell proliferation. (A) Box-plot quantification of vascular cell number (all cells in the stele, excluding the pericycle). Asterisks indicate statistically significant difference from Col-0 after pairwise t-test with ***p < 0.001. (B) Response of root length to depletion and exogenous addition of brassinosteroids in Col-0 and *rlp44^cnu2^*. Bars denote mean root length ± SD, n = 14-26. Asterisks denote statistically significant differences after two-way ANOVA and Tukey’s post hoc test between Col-0 and *rlp44^cnu2^* with **p < 0.01 and *** p<0.001. (C) Response of vascular cell number to depletion (PPZ) and exogenous addition of brassinosteroids (epi-brassinolide, BL) in Col-0 and *rlp44^cnu2^*. Asterisks indicate statistically significant differences from control conditions as determined by two-tailed t test with **p < 0.01 and *** p<0.001.

### RLP44 controls xylem cell fate by promoting phytosulfokine signalling

The results described so far suggested that the maintenance of procambial cell identity in the root requires the presence of both BRI1 and RLP44. In addition, RLP44 seems to act downstream of BRI1 in controlling vascular cell fate. As it could be ruled out that RLP44 exerts its effect on xylem differentiation through BR signalling, we reasoned that in the absence of a kinase domain, RLP44 is required to interact with and influence the activity of another signalling component which, in turn, controls xylem cell fate. Interestingly, besides BRI1 and its close homologues BRL1, BRL2, and BRL3, the LRR X clade of RLKs harbours the receptors for the peptide growth factor PSK, PSKR1 and 2 (16). As PSK signalling has also been implicated in promoting the trans-differentiation of *Zinnia elegans* mesophyll cells into tracheary elements (31, 47) and depends on functional BR signalling (29), we tested the association of RLP44 with PSKR1. Co-immunoprecipitation experiments in *N. benthamiana* showed that *PSKR1-GFP* (33) was present in RLP44-RFP immunoprecipitates (Fig. 5A). In addition, FRET-FLIM analysis showed a pronounced reduction in fluorescence lifetime when *PSKR1-GFP* was co-expressed with RLP44-RFP, suggesting a direct interaction (Fig. 5B) that was not affected by exogenous application of PSK (Fig. S7A and B). Moreover, exogenous application of PSK peptide reverted *rlp44* xylem back to the stereotypical wildtype pattern of mostly two protoxylem and three metaxylem cells in one axis (Fig. 5C). Further supporting a role of PSK signalling in the control of xylem cell fate, the *pskr1-3 pskr2-1* double mutant (48) showed an increased number of xylem cells reminiscent of the *rlp44* mutant (Fig. 6D and E). A similar phenotype was observed in the double mutant of *pskr1-3* and the related RLK *psy1r1*, as well as in the *tpst-1* mutant, impaired in the biosynthesis of PSK and other sulfated peptides (Fig. 5D and E) (28, 49). We then asked how RLP44 might promote PSK signalling. As RLP44 is a direct interaction partner of both PSKR1 and its co-receptor BAK1, we assessed whether presence of RLP44 could increase association of receptor and co-receptor. Indeed, more BAK1 was detected in immunoprecipitates of PSKR1-GFP expressed in *N. benthamiana* when RLP44-RFP was co-expressed (Fig. 5F), suggesting that RLP44 might act as a scaffold in the complex. Supporting these results, BAK1 levels in immunoprecipitates of PSKR1-GFP were reduced in the *rlp44^cnu2^* mutant (Fig. 5G). Notably, presence of PSKR1-GFP had a negative effect on the amount of RLP44 in immunoprecipitates of BRI1-RFP (Fig. S7C), suggesting that the two pathways might compete for RLP44. Taken together, our data suggest that RLP44 interacts with, and stabilizes a complex between, PSKR1 and BAK1 to promote PSK signalling, which, in turn, suppresses the progression from procambial to xylem identity (Fig. 5H).

**Fig. 5.**
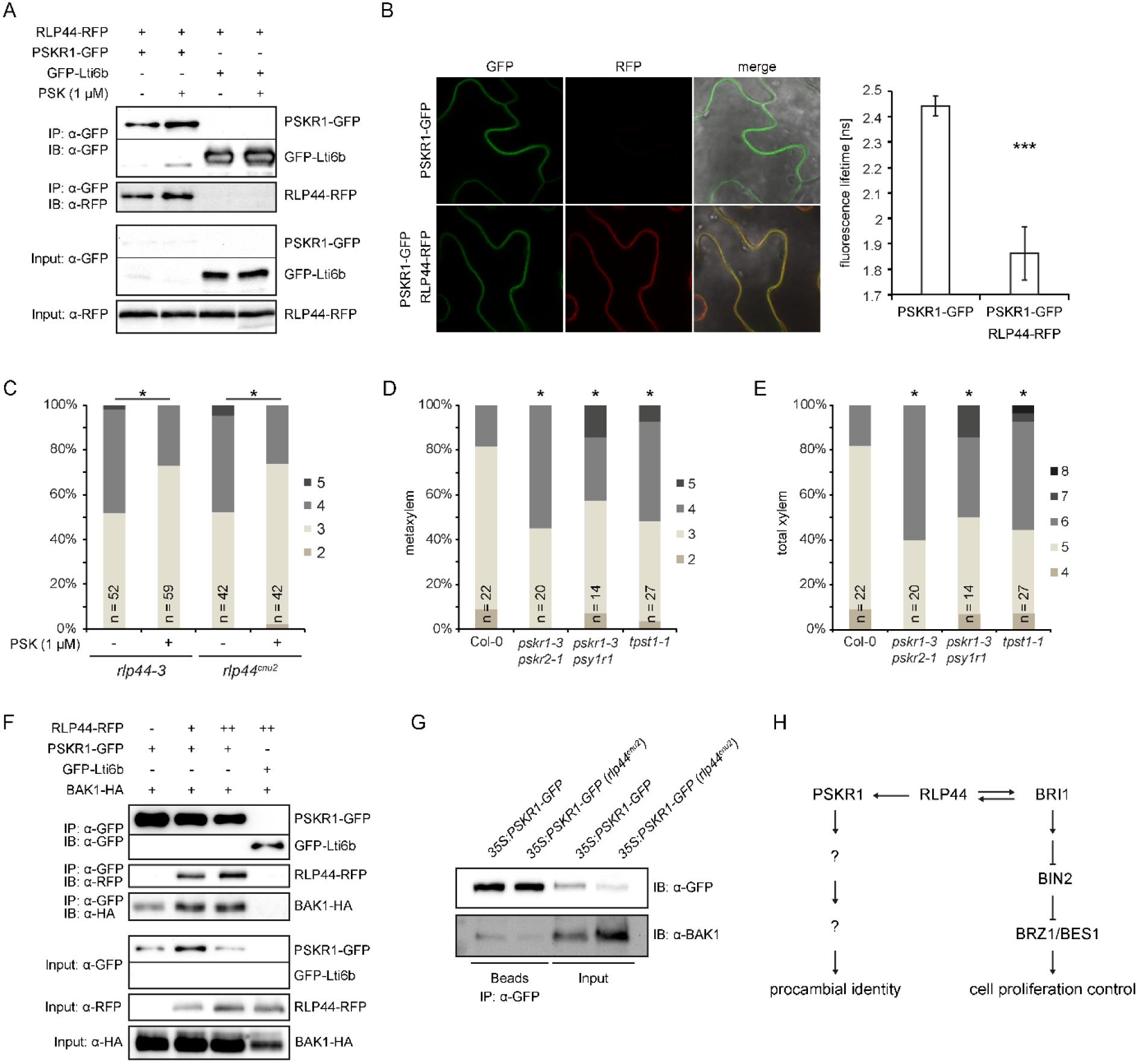
RLP44 interacts with PSKR1 to promote PSK signalling and procambial identity. (A) Co-immunoprecipitation after transient expression in *Nicotiana benthamiana* leaves demonstrates presence of RLP44-RFP in PSKR1-GFP immunoprecipitates, in contrast to immunoprecipitates of the Lti6b-GFP control. (B) FRET-FLIM analysis of the PSKR1-GFP/RLP44-RFP interaction in *Nicotiana benthamiana* leaves. Bars denote average of 9 measurements ± SD. Asterisks indicate statistically significant difference from PSKR1-GFP according to pairwise t-test (***p < 0.001). (C) Application of PSK peptide rescues the ectopic xylem phenotype of *rlp44^cnu2^*. Asterisks indicate statistically significant difference according to Mann-Whitney U-test (*p < 0.05). (D and E) Quantification of metaxylem (D) and total xylem (E) cell number in Col-0 and PSK signalling-related mutants. Asterisks indicate statistically significant difference from Col-0 based on Dunn’s post-hoc test with Benjamini-Hochberg correction after Kruskal-Wallis modified U-test (*p < 0.05). (F) Co-immunoprecipitation analysis after transient expression in *Nicotiana benthamiana* leaves demonstrates increased amount of BAK1-HA in PSKR1-GFP immunoprecipitates in the presence of RLP44-RFP. RLP44 levels were adjusted through increasing the density of Agrobacteria (denoted by + or). (G) BAK1-HA is decreased in immunoprecipitates of PSKR1-GFP in the *rlp44^cnu2^* background. (H) Model of RLP44-mediated integration of PSK and BR signalling.

## Discussion

### RLP44 controls vascular cell fate in a BRI1-dependent manner

Cell fate determination in plants mainly relies on positional information provided by perception of hormone gradients and non-cell autonomous factors (50-52). The expanded family of plant RLK proteins and their ligands play central roles in intercellular communication, cell identity maintenance, and the regulation of cell expansion and proliferation (53). Currently, our view of these pathways is evolving to appreciate the extensive cross-talk and interdependence of diverse signalling pathways (54). The response to BRs, mediated by what is probably the best-characterized plant signalling cascade (11), is found to be integrated with a growing number of other pathways at two cross-talk “hot spots”, namely the GSK3-like kinase BIN2 (55-57) and the BR-responsive transcription factors BZR1 and BES1 (58-61). Here, we report that BR and PSK signalling are coupled at the level of the plasma membrane receptors through RLP44 and that this signalling module is required to control xylem cell fate. Our genetic and biochemical analyses support a scenario where PSK signalling strength is quantitatively controlled by RLP44, which itself is dependent on the presence of BRI1. While we do not rule out post-translational control of RLP44 by BRI1, for example through phosphorylation of the cytoplasmic domain or because the presence of BRI1 is required for correct receptor complex assembly, this dependency is at least partially based on transcriptional regulation. The *bri1-null* loss-of-function mutant, but not hypomorphs or the BR biosynthetic mutant *cpd*, showed reduced RLP44 expression. Consistent with this, uncoupling RLP44 from its native transcriptional regulation by constitutive expression could rescue the *bri1-null* xylem phenotype. Conversely, the novel *bri1* loss-of-function phenotype of ectopic xylem in procambial position is unrelated to BR-signalling. In line with this, only limited overlap of differentially expressed genes in *bri1-116* and the morphologically similar BR biosynthetic mutant *dwf4* was observed, potentially indicating additional brassinosteroid-independent functions of BRI1 (21).

More work will be needed to mechanistically understand the interaction between RLP44 and RLKs. External application of PSK (this study) or BR (34) ligands had no influence on the association between RLP44 and PSKR1 or BRI1, respectively, in line with ligand-independence of many, but not all, RLP-RLK interactions (62, 63). BRI1 and PSKR1/2 belong to the LRR X subfamily of LRR-RLKs (16) and share the requirement for interaction with SERK co-receptors to form an active, heteromeric signalling complex (32, 33, 64). With the exception of the ligand binding domain, the SERK-bound BRI1 and PSKR1 complex structures are very similar (32, 64), therefore it is conceivable that RLP44 binds both receptors through the same mechanism. These results are in line with the emerging theme of dynamic, promiscuous, and flexible interactions of plasma membrane proteins to integrate signalling information and fine-tune cellular responses to external cues (65-67). Interestingly, the mechanism by which RLPs influence signalling seems to differ widely, ranging from direct participation in ligand binding (62, 68), to the control of signalling specificity through blocking access of RLK ligands (68), to the guarding of extracellular proteins targeted by pathogens (69). Here, we propose a scaffolding function of RLP44 for the interaction between PSKR1 and its co-receptor BAK1, expanding the mechanistic diversity of RLPs. We have previously reported that RLP44 is involved in the response to cell wall state; along those lines, it will be interesting to see whether cues from the cell wall are able to influence the balance between the two known roles of RLP44 by shifting its interaction with BRI1 or PSKR1.

### PSK signalling likely promotes procambial identity

Alongside classical plant hormones, signalling peptides have been revealed to play major roles in plant development and stress responses (70, 71). The sulfated pentapeptide PSK has been implicated in a number of diverse processes (24, 71). Here, we propose that PSK signalling controls xylem cell fate through promoting the maintenance of procambial identity. A number of observations support this hypothesis. First, PSK treatment rescued the ectopic xylem phenotype in *rlp44* mutants. Second, PSK-related mutants showed increased xylem differentiation in procambial position and PSK genes are co-expressed with RLP44 in procambial cells (Fig. S7D)(38). Third, PSK expression is transiently increased prior to the acquisition of a procambial intermediate state by cells trans-differentiating into tracheary elements (31, 72), which could explain why PSK promotes tracheary element formation in *Z. elegans* only when applied early to the cell culture (31, 47). Finally, PSK signalling promotes callus growth and longevity, in line with a role in the maintenance of cell identity (27). Accordingly, it has been proposed that PSK signalling maintains the responsiveness to intrinsic and extrinsic cues to tune proliferation or differentiation (27). However, it is unclear how PSK signalling affects cellular behaviour, in part due to a lack of knowledge about potential downstream targets. It will be interesting to see how the BRI1-RLP44-PSK signalling module described here integrates with the fundamental patterning processes and the gene regulatory networks controlling xylem differentiation (2, 7).

### The role of BR signalling in vascular development

It has long been described that BR signalling plays an important role in the development of vascular tissue (13, 14). In addition, it has been reported that BR signalling is kept at low levels in procambial cells of leaf and hypocotyl to prevent their differentiation into xylem cells (57). Our results suggest that in the primary xylem of the root, BR signalling only plays a minor role in controlling differentiation, in marked contrast to the strong patterning defects of BR signalling and biosynthetic mutants in the shoot (14). Conversely, at least in the root, presence of BRI1 has a negative effect on xylem cell fate through RLP44 and PSK signalling-mediated maintenance of procambial identity. Therefore, our results identify a novel role of BRI1 in root development which is independent of its role as BR receptor.

## Materials and Methods

Details of materials and methods are provided in the Supplemental Information file. Mutants and transgenic lines used in this study are listed in Table S1.

## Acknowledgements

The authors would like to thank Sigal Savaldi-Goldstein for the *bri* triple mutant, Michael Hothorn for antisera against BRI1 and BAK1, and Karin Schumacher for antiserum against RFP and for critical reading of the manuscript. Research in our laboratories was supported by the German Research Foundation (DFG) with grants to SW (WO 1660/6-1) and to KH (HA 2146/22-1; CRC 1101-D02). SW is supported by the DFG through the Emmy Noether Programme (WO 1660/2-1).

